# Optimized Mucosal MVA Prime/ Soluble gp120 Boost Vaccination Regimen Induces Similar Antibody Responses as an Intramuscular Regimen

**DOI:** 10.1101/573394

**Authors:** Dorothy I. Jones, Justin J. Pollara, Brandi T. Johnson-Weaver, Celia C. LaBranche, David C. Montefiori, David J. Pickup, Sallie R. Permar, Soman N. Abraham, Massimo Maddaloni, David W. Pascual, Herman F. Staats

**Affiliations:** Departments of Pathology, Duke University School of Medicine, Durham, NC 27710; Departments of Surgery, Duke University School of Medicine, Durham, NC 27710; Departments of Molecular Genetics and Microbiology, Duke University School of Medicine, Durham, NC 27710; Departments of Pediatrics, Duke University School of Medicine, Durham, NC 27710; Departments of Immunology, Duke University School of Medicine, Durham, NC 27710; Departments of Medicine, Duke University School of Medicine, Durham, NC 27710; Departments of the Human Vaccine Institute, Duke University School of Medicine, Durham, NC 27710; Department of Infectious Diseases & Immunology, College of Veterinary Medicine, University of Florida, Gainesville, FL 32608

## Abstract

The benefits of mucosal vaccines over injected vaccines are difficult to ascertain since mucosally administered vaccines often induce serum antibody responses of lower magnitude than those induced by injected vaccines. This study aimed to determine if mucosal vaccination using a modified vaccinia Ankara expressing HIV-1 gp120 (MVA-g120) prime and HIV-1 gp120 protein boost could be optimized to induce serum antibody responses similar to those induced by an intramuscularly (IM) administered MVA prime/gp120 boost to allow comparison of an IM immunization regimen to a mucosal vaccination regimen for their ability to protect against a low dose rectal SHIV challenge while inducing similar serum anti-HIV-1 antibody responses. A 3-fold higher antigen dose was required for intranasal (IN) immunization with gp120 to induce serum anti-gp120 IgG responses not significantly different than those induced by IM immunization. Gp120 fused to the Adenovirus type 2 fiber binding domain (gp120-Ad2F), a mucosal targeting ligand, exhibited enhanced IN immunogenicity when compared to gp120 alone. MVA-gp120 was more immunogenic after IN delivery than gastric or rectal delivery, although serum antibodies induced by IN immunization were lower than those induced by intramuscular immunization. Using these optimized vaccines, an IN MVA-gp120 prime, combined IM (gp120) and IN (gp120-Ad2F) boost regimen (IN/IM+IN) induced serum anti-gp120 antibody titers similar to those induced by the intramuscular prime/boost regimen (IM/IM) in rabbits and non-human primates. Despite the induction of similar systemic anti-HIV-1 antibody responses, neither the IM/IM nor the IN/IM+IN regimen induced elevated anti-HIV-1 mucosal IgA responses nor protected against a repeated low-dose rectal SHIV challenge. These results demonstrate that immunization regimens utilizing the IN route are able to induce serum antigen-specific antibody responses similar to those induced by systemic immunization

**IMPORTANCE:** Mucosal vaccination is proposed as a method of immunization able to induce protection against mucosal pathogens that is superior to protection provided by parenteral immunization. However, mucosal vaccination often induces serum antigen-specific immune responses of lower magnitude than those induced by parenteral immunization, making the comparison of mucosal and parenteral immunization difficult. We identified vaccine parameters that allowed an immunization regimen consisting of an IN prime followed with boosters administered by both IN and IM routes to induce serum antibody responses similar to those induced by IM prime/boost vaccination. Additional studies are needed to determine the potential benefit of mucosal immunization for HIV-1 and other mucosally-transmitted pathogens.

## INTRODUCTION

An HIV vaccine would be of significant benefit for ending the HIV/AIDS epidemic (1). To date, the only HIV vaccine clinical trial to demonstrate protective efficacy was the pox virus prime/recombinant gp120-boost RV144 trial (NCT00223080) (2). Since more than 90% of HIV infections are transmitted across mucosal tissues, with sexual transmission being the most common route (3), enhancing mucosal immunity with the use of mucosal immunization may enhance protection against sexual transmission of HIV(4). The benefit of mucosal immunity for protection against mucosal transmission of SHIV/SIV has been demonstrated using passive and active immunization. For example, passive transfer of the combination of a systemically administered anti-HIV IgG1 and a mucosally administered anti-HIV dimeric IgA2 provided complete protection against a high-dose rectal SHIV challenge while transfer of the anti-HIV IgG1 alone was not protective (5), suggesting that the combination of systemic and mucosal anti-HIV IgG and IgA, respectively, may be required for maximum protection. Another recent study demonstrated that systemic (intramuscular) and mucosal (aerosol) immunization with the same vaccine induced equivalent levels of protection against SIV challenge(6). However, evaluation of vaccine-induced immune responses determined that protection induced by intramuscular immunization was associated with serum IgG-mediated immune responses while protection induced by aerosol immunization was associated with serum IgA-mediated immune responses(6) Therefore, optimizing both systemic and mucosal immunization regimens may be required to provide consistent protection against mucosal HIV transmission(7).

HIV-1 vaccine regimens utilizing a mucosal route of vaccination have been described in the literature but evaluating the potency of a mucosally-administered vaccine by comparison to a similar vaccine delivered parenterally is often not discussed. For example, mucosal immunization often fails to induce antigen-specific serum IgG responses comparable to those induced by parenteral immunization with the same antigen(8, 9). For example, our previous study that compared IM MVA priming with IM gp120 boosting (IM/IM) to IM MVA priming with IN gp120 boosting (IM/IN) demonstrated unique immune responses dependent on the route of immunization used(8). While the IM/IM and IM/IN vaccine regimens both induced similar levels of anti-HIV binding antibodies, the IM/IN vaccine regimen enhanced the induction of breast milk anti-HIV IgA responses while it did induce HIV-1 neutralizing antibody responses comparable to those induced by the IM/IM vaccine regimen, indicating that IM/IM systemic immunization was more effective than IM/IN mucosal vaccination at eliciting functional systemic antibody responses. Others have reported that IN immunization of humans with 100 µg of HIV-1 gp140 was not immunogenic while IM immunization with 20 or 100 µg of the same antigen induced elevated serum anti-gp140 IgG responses(9). This observation highlights the difficulty in comparing the immunogenicity of the same HIV-1 antigen dose delivered parenterally vs mucosally. If the mucosal immunization strategy is not immunogenic, it is impossible to evaluate correlates of protection for the mucosal immunization strategy as compared to the parenteral immunization strategy, as described above (8). We therefore propose that one criteria to identify an optimized mucosal immunization regimen is the expectation that the mucosal immunization regimen induce systemic anti-HIV binding and functional antibody responses, such as virus neutralization or antibody-dependent cell-mediated cytotoxicity (ADCC), comparable to those induced by systemic immunization, while utilizing a mucosal route of immunization that may enhance the induction of protective serum(6) or mucosal (5) anti-HIV IgA responses.

A major impediment to developing any mucosal vaccine is the lack of an approved mucosal vaccine adjuvant. Of all the mucosal vaccine adjuvants tested to date, cholera toxin (CT) has been one of the most efficacious and has been used as a “gold standard” for mucosal adjuvant efficacy(10). However, safety concerns, including nerve toxicity, will likely prevent it from being used in humans as a nasal vaccine adjuvant(10). Monophosphoryl Lipid A (MPL) is a toll-like receptor 4 (TLR4) agonist that is approved for use in injected vaccines when it is combined with alum and clinical trials indicate MPL is safe for use as a nasal vaccine adjuvant (11).. Since mucosal vaccines need to be both safe and efficacious, newer compounds have been investigated as mucosal vaccine adjuvants including mast cell activating compounds and cationic peptides, which can bridge innate and adaptive immune responses (12, 13). Compound 48/80 is a mast cell activating compound with adjuvant activity capable of inducing antibody responses similar to CT-adjuvanted vaccines, when used as an adjuvant for mucosal or systemic immunization (14-17). However, C48/80, represents a mixture of polymer species (18), and therefore it may not be an ideal adjuvant as regulatory agencies would most likely require a single active compound to provide the described adjuvant activity. To identify a potent adjuvant for nasally administered HIV-1 gp120, we compared the ability of several adjuvants— CT, MPL, C48/80, as well as mastoparan 7 (M7), a cationic antimicrobial mast cell activating peptide (19, 20)— to enhance serum anti-gp120 antibody responses.

In addition to immunostimulatory adjuvants, adjuvants that enhance vaccine retention at mucosal sites may be beneficial. Several studies have demonstrated that increased nasal clearance correlates with diminished immunogenicity of nasal vaccines (21-25). To enhance nasal vaccine retention, we tested a fused protein consisting of gp120 and the adenovirus type 2 fiber protein (Ad2F). Ad2F binds the coxsackie and adenovirus receptor (CAR) expressed on epithelial and endothelial cells, and thus can act as a mucosal targeting ligand to enhance antigen binding to the epithelial surfaces in the nasal cavity(26). In addition to enhancing vaccine retention, Ad2F binding can induce the release of proinflammatory cytokines and chemokines believed to impart a localized adjuvant effect (27, 28). The enhanced immunogenicity of fusion proteins consisting of botulinum neurotoxin vaccine antigens and Ad2F have previously been reported(17, 26, 29). However, the ability of Ad2F to enhance the nasal immunogenicity of HIV-1 gp120 has not been evaluated. Here we test a fusion protein composed of HIV-1 gp120 and Ad2F to determine if the addition of Ad2F enhances the nasal immunogenicity of HIV-1 gp120.

Viral vectored vaccines are another vaccine/adjuvant system that may enhance the immunogenicity of mucosal vaccine regimens. Viruses that can infect the host through mucosal tissues, such as modified vaccinia Ankara (MVA), have mechanisms to overcome the innate immune defenses at mucosal sites and may be more easily adapted to mucosal administration than subunit vaccines. Injected MVA vectored vaccines have been used in several HIV vaccine clinical trials, but these have shown various degrees of efficacy (30-36). Strategies that enhance insert-specific immune responses and/or minimize vector-specific responses may enhance efficacy of MVA vaccine vectors. Thus, replication-defective MVA viral vectors may offer advantages over conventional MVA vectors. For example, MVA vectors lacking the ability to express late genes, immunomodulatory proteins or lacking the ability to replicate may be induce elevated insert-specific immune responses when compared to conventional MVA vectors (37-39) (40-42).. In this study, we evaluated the immunogenicity of traditional MVA and anMVA vector lacking the replication gene (udg), both expressing HIV-1 gp120,when delivered intramuscularly or mucosally (intranasally, intragastric or rectally).

Here we identify an HIV-1 vaccination regimen that utilizes a mucosal route of immunization that induces serum gp120-specific IgG titers similar to those induced by an injected vaccine regimen. Using the adjuvant systems discussed above, we optimized a vaccination regimen consisting of an MVA expressing HIV-1 gp120 prime followed by adjuvanted gp120 booster immunizations. Rabbits were used as a small animal model for the optimization of mucosal vaccines. We demonstrate the translatability of our optimized mucosal vaccination regimen from rabbits to nonhuman primates (NHPs) by comparing the efficacy of the optimized mucosal vaccination regimen to the systemic regimen in rhesus macaques. Since the NHP model allows us to begin investigating the potential protective benefit to including mucosal vaccines in an HIV vaccination regimen, we rectally challenged both the mucosally and systemically vaccinated macaques with a heterologous tier 2 SHIV.

## METHODS

### MVA vaccines

For experiment “MVA 1” in **Table 1**, MVA expressing HIV-1 C.1086 gp140 was used. MVA for all other vaccinations expressed HIV-1 C.1086 gp120. Gp120 or gp140 was placed under the control of a synthetic promoter capable of high levels of transcription in both early and late stages of viral replication and inserted between two essential genes as previously described (8, 43). HIV protein expression was confirmed by Western blot. Replication-defective MVA was produced by deleting uracil-DNA-glycosylase (*udg*), which is essential for viral replication (44). See Supplemental Information for more details on the construction of the various MVA vectors.

**Table 1.**
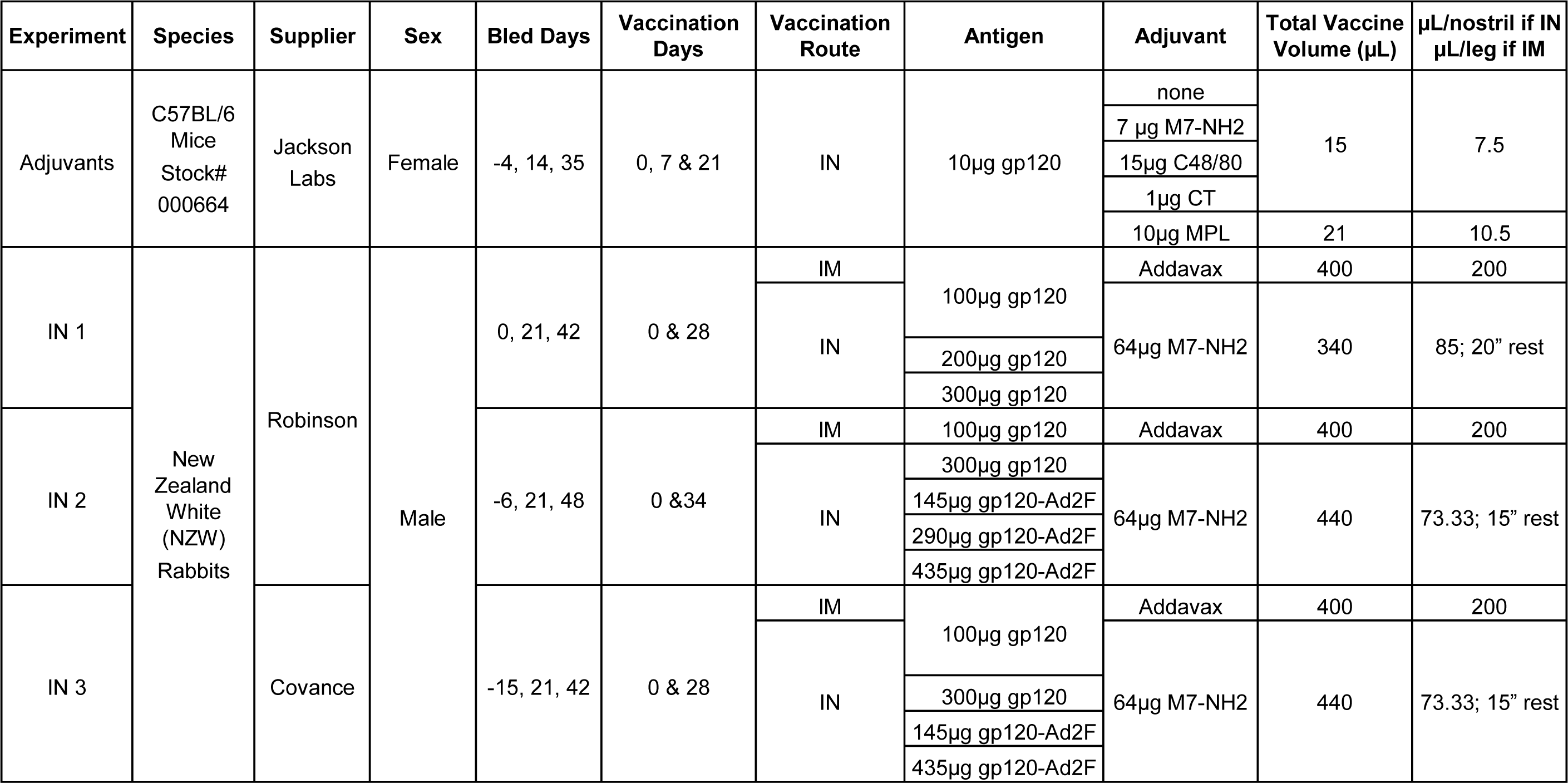
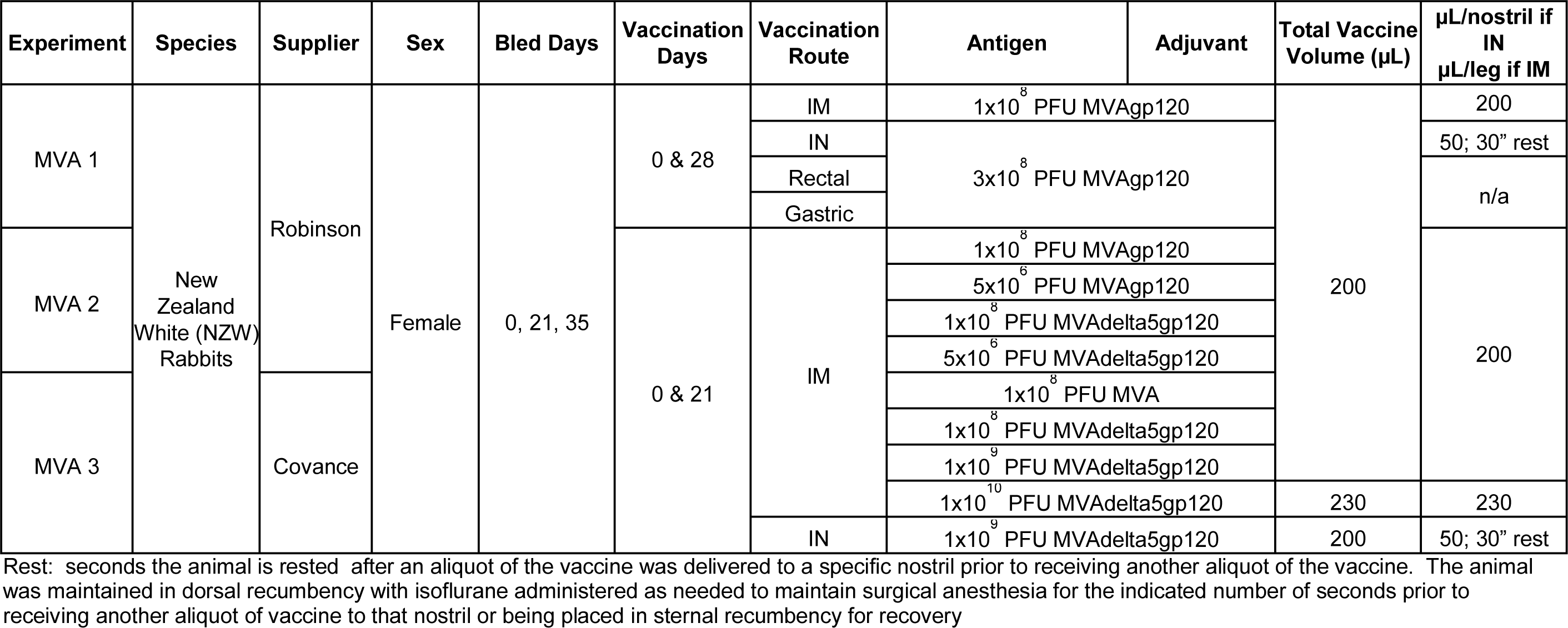
Animal characteristics and vaccine specifications for optimization experiments in mice and rabbits.

### Protein Vaccines and Adjuvants

The antigen used for booster immunizations consisted of either C.1086 delta7gp120K160N (gp120) (45, 46) or a fusion protein between C.1086 delta7gp120K160N and the Ad2F adhesin (gp120-Ad2F) using the method to that previously described (26). Briefly, the C-terminal region of the Ad2F, starting from G378 to E582, was used as the transporter/targeting domain. This region encompasses 1) a short stretch of amino acids rich in Gly (4 of 15); 2) a trimerizing domain; and 3) a globular domain, commonly referred to as the “knob,” which is important for interacting with the coxsackievirus/adenovirus receptor on the cell surface. The C.1086 delta7gp120K160N ATG codon was embedded in an ideal Kozak’s sequence, cloned upstream of the Ad2F transporter, and the carboxy-terminus of the fusion contained a His-tag to allow for protein purification. At both ends of the synthetic gene, two XbaI sites allowed for excision and cloning into the mammalian expression vector pcDNA3.1+ (Invitrogen). Orientation of the clones was determined by restriction analysis and confirmed by DNA sequencing. The resulting construct named pGnMM9 was used for protein expression by Paragon (http://paragonbioservices.com/). Briefly, 293T cells were transfected with a synthetic gene encoding the fusion protein gp120-Ad2F. Secreted gp120-Ad2F was affinity purified by lectin affinity columns, and purity was assessed by SDS-page and western blot.

Intramuscular vaccines adjuvanted with Addavax (Invivogen, cat #vac-adx-10) were mixed 1:1 with antigen for total vaccine volume. Intranasal vaccines tested in rabbits and macaques were adjuvanted with 64 µg of the antimicrobial, mast cell activating peptide Mastoparan-7 (INLKALAALAKALL-NH2; CPC Scientific, Sunnyvale, CA). In mice, intranasal vaccines were adjuvanted with 7 µg M7-NH2 (CPC), 15 µg compound 48/40 (Sigma), 1 µg cholera toxin (Enzo Life Sciences), or 10 µg MPL (Enzo Life Sciences). USP grade 0.9% saline (APP Pharmaceuticals) was added as needed to create total vaccine volume, which is listed in **Table 1.**

### Animal husbandry and procedures

All experimental procedures were performed in accordance with IACUC policies. Information on husbandry, health events, anesthetic protocols and other detailed information necessary to meet the ARRIVE guidelines (47) is available in **Supplemental Information**. A summary describing the number of experiments conducted, animal characteristics, as well as the vaccine formulation are provided in **Tables 1 and 2**.

**Table 2.**
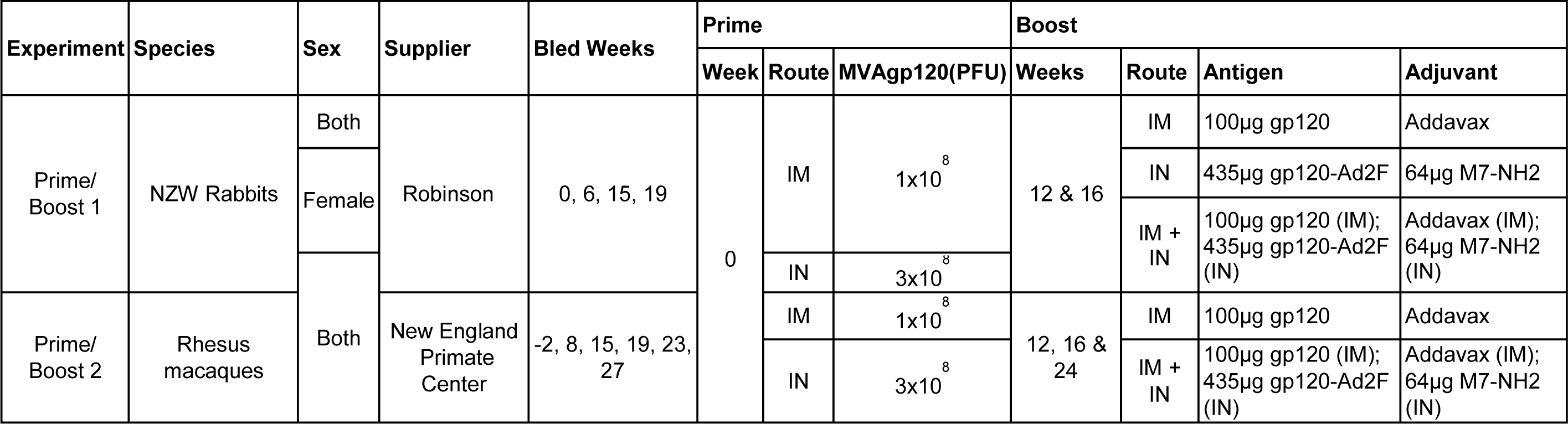
Animal characteristics and vaccine specifications for prime/boost experiments.

### Mouse vaccination and sample collection

Female C57BL/6J (Jackson Labs; 8 weeks old) were vaccinated by the intranasal (IN) route on day 0, 7, and 21. Serum was collected on day −4, 14, and 35. Vaginal lavage was collected as previously described (48, 49) on day −4 and 35.

### Rabbit vaccination and sample collection

New Zealand White rabbits greater than 2kg were obtained from Robinson Services (Mocksville, NC) or Covance Research Products (Denver, PA). For vaccine optimization experiments, two vaccines were administered 3-5 weeks apart. Rabbits were bled prior to the first vaccination, 3 weeks after the first vaccination, and 1-2 weeks after the second vaccination. For prime/boost experiments, rabbits received the MVA-gp120 prime on week 0 and gp120 protein boosts on weeks 12 and 16. They were bled on weeks 0, 6, 15, and 19.

### NHP vaccination and sample collection

Sixteen 1-3 year old rhesus macaques were obtained from the New England Primate Research Center. They received the MVA-gp120 prime on week 0 and gp120 protein boosts on weeks 12, 16, and 24. They were bled on weeks −2, 8, 15, 19, 23, and 27. SHIV challenges began on week 30.

### SHIV Challenge

Rectal challenges with a heterologous tier-2 SHIV-1157QNE Y173H (50, 51) (2014 stock, provided by Sampa Santra) were administered weekly for up to 10 weeks or until infection was detected.. For the first four challenges virus stock was diluted 1:10,000. After the fourth challenge, virus stock dilution decreased ten-fold every other challenge (challenge 5 & 6 1:1,000; challenge 7 & 8 1:100; challenge 9 & 10 1:10). All challenges were diluted with 0.5% human serum albumin (30% solution, Calbiochem) in PBS and administered in a 1mL volume. Four unvaccinated macaques were used as naïve controls.

### ELISA

ELISA assays were performed as described (52) with the following exceptions. Coating antigen was C.1086 K160N gp120 or V1V2 tags (NIH AIDS reagent program, Cat#12568) at 2 µg/mL. To detect antibody responses to the viral vector in animals immunized with MVA, the vaccinia virus proteins L1 and B5 (BEI cat# NR-2622, NR-545, NR-2625, NR-546) were used as coating antigens at 2 µg/mL. Serum samples were tested starting at a 1:32 dilution while mucosal samples were tested starting at a 1:16 dilution. Serial 2-fold dilutions were performed. If any individual animal had titers >2^27^ at a given time point, samples from all animals in the experiment were retested using serial 3-fold dilutions. Murine antibodies were detected using alkaline phosphatase conjugated goat anti-mouse IgG or IgA (Southern Biotech, Cat #1030-04 and 1040-04). Rabbit antibodies were detected using alkaline phosphatase conjugated goat anti-rabbit IgG (Southern Biotech, Cat. #4030-04) or IgA (Novus, Cat. #NB7167). For NHP studies, the secondary antibodies used were anti-NHP IgG and anti-NHP IgA (Rockland, cat. #617-105-012, 617-105-006). Endpoint titers are defined as the last immune sample dilution with an ELISA raw data value three-fold higher than that animal’s pre-immune sample at the same dilution. Since the mice are inbred, the average of two pre-immune samples was used instead of each individual’s. Immune samples that were not three-fold higher than the naïve sample at the starting dilution were assigned the value of the one dilution lower than the starting dilution.

### Antibody Dependent Cellular Cytotoxicity

ADCC activity was measured using the GranToxiLux (GTL) assay as previously described (53). A clonal isolate of the CEM.NKR_CCR5_ CD4^+^ T cell line (NIH AIDS Reagent Program, Division of AIDS, NIAID, NIH: from Dr. Alexandra Trkola (54)) was used as the source of target cells after coating with the recombinant HIV-1 gp120 protein used as the boost component of the vaccine regimen (HIV-1 isolate C.1086). Effector cells were human PBMC obtained from a HIV-seronegative donor heterozygous for Fcγ Receptor IIIa at position 158 (158F/V) (55, 56). PBMC and target cells were combined at an effector cell to target cell ratio of 30:1. Samples were tested after four-fold serial dilutions starting at 1:100. ADCC activity was measured as the proportion of cells positive for proteolytically active granzyme B (GzB) out of the total viable target cell population (%GzB activity), after subtracting the background activity observed in wells containing effector and target cells in the absence of plasma. ADCC endpoint titers were determined by interpolating the dilutions of plasma that intercepted the previously established positive cutoff for this assay (8% GzB activity) using GraphPad Prism, version 7.0b software (GraphPad Software, Inc.), and are reported as reciprocal dilution.

### Neutralization

Neutralizing antibody activity was measured in 96-well culture plates by using Tat-regulated luciferase (Luc) reporter gene expression to quantify reductions in virus infection in TZM-bl cells. nTZM-bl cells were obtained from the NIH AIDS Research and Reference Reagent Program, as contributed by John Kappes and Xiaoyun Wu. Assays were performed with HIV-1 Env-pseudotyped viruses as described previously (57, 58). Test samples were diluted over a range of 1:20 to 1:43740 in cell culture medium and pre-incubated with virus (∼150,000 relative light unit equivalents) for 1 hr at 37 oC before addition of cells. Following a 48 hr incubation, cells were lysed and Luc activity determined using a microtiter plate luminometer and BriteLite Plus Reagent (Perkin Elmer). Neutralization titers are the sample dilution at which relative luminescence units (RLU) were reduced by 50% compared to RLU in virus control wells after subtraction of background RLU in cell control wells. Serum samples were heat-inactivated at 56oC for 45 min (rabbits) or 30 min (NHP) prior to assay.

### Viral Load Quantification

qPCR was used to quantify SHIV RNA present in serum collected one week after each challenge. Viral RNA measurements were performed by the Immunology Virology Quality Assessment Center Laboratory Shared Resource, Duke Human Vaccine Institute, Durham NC as previously described (59). The lower limit of detection was 250 copies per mL.

### Statistical methods

Mouse and rabbit data was analyzed by Kruskal-Wallis with Dunn’s multiple comparisons test. NHP ELISA data was analyzed by Mann-Whitney. Graph preparation and all non-parametric comparisons were performed in Graphpad Prism (https://www.graphpad.com/;La Jolla, CA). Values from individual animals as well as the geometric mean titer is shown with the geometric mean of each group depicted on the graph as the horizontal black bar unless otherwise stated.

## RESULTS

### Evaluation of non-toxin adjuvants for use with nasally-administered HIV-1 gp120

The lack of an approved mucosal vaccine adjuvant is a major deterrent to mucosal vaccine development. To determine if a non-toxin adjuvant could provide adjuvant activity similar or superior to CT, we compared MPL, C48/80, and M7 to the “gold standard” of CT for the ability to enhance serum antibody responses to HIV-1 gp120 when IN administered to C57BL/6 mice. On day 14, all mice (5/5) that were vaccinated with gp120 + C48/80 or mastoparan 7 (M7) had detectable serum anti-gp120 IgG response compared to 4/5 mice vaccinated with gp120 + MPL, 3/5 mice vaccinated with gp120 + CT, and 2/5 mice that received gp120 alone (**Table 3**). Only the mice that received gp120 + M7 had serum anti-gp120 IgG titers significantly (p=0.005) different than the mice that received gp120 alone (GMT 1:4,782,969 vs 1:33.63 respectively).

**Table 3.**
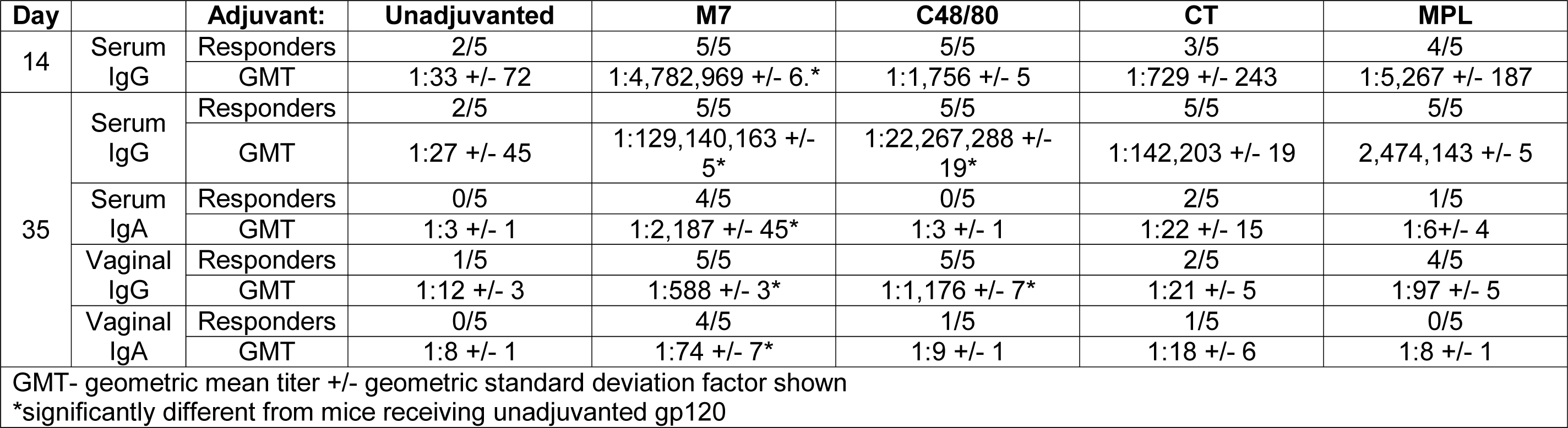
Murine gp120-specific antibody responses after intranasal vaccination.

| Day |  | Adjuvant: | Unadjuvanted | M7 | C48/80 | CT | MPL |
| --- | --- | --- | --- | --- | --- | --- | --- |
| 14 | Serum IgG | Responders | 2/5 | 5/5 | 5/5 | 3/5 | 4/5 |
|  |  | GMT | 1:33 +/- 72 | 1:4,782,969 +/- 6.* | 1:1,756 +/- 5 | 1:729 +/- 243 | 1:5,267 +/- 187 |
| 35 | Serum IgG | Responders | 2/5 | 5/5 | 5/5 | 5/5 | 5/5 |
|  |  | GMT | 1:27 +/- 45 | 1:129,140,163 +/- 5* | 1:22,267,288 +/- 19* | 1:142,203 +/- 19 | 2,474,143 +/- 5 |
|  | Serum IgA | Responders | 0/5 | 4/5 | 0/5 | 2/5 | 1/5 |
|  |  | GMT | 1:3 +/- 1 | 1:2,187 +/- 45* | 1:3 +/- 1 | 1:22 +/- 15 | 1:6+/- 4 |
|  | Vaginal IgG | Responders | 1/5 | 5/5 | 5/5 | 2/5 | 4/5 |
|  |  | GMT | 1:12 +/- 3 | 1:588 +/- 3* | 1:1,176 +/- 7* | 1:21 +/- 5 | 1:97 +/- 5 |
|  | Vaginal IgA | Responders | 0/5 | 4/5 | 1/5 | 1/5 | 0/5 |
|  |  | GMT | 1:8 +/- 1 | 1:74 +/- 7* | 1:9 +/- 1 | 1:18 +/- 6 | 1:8 +/- 1 |
| GMT- geometric mean titer +/- geometric standard deviation factor shown |  |  |  |  |  |  |  |
| *significantly different from mice receiving unadjuvanted gp120 |  |  |  |  |  |  |  |

By day 35, all mice (5/5) that were vaccinated with an adjuvanted vaccine developed detectable serum anti-gp120 IgG titers compared to only 2 of 5 mice vaccinated with gp120 alone. However, only gp120 adjuvanted with C48/80 or M7 induced serum anti-gp120 IgG titers significantly greater than the anti-gp120 IgG induced by the unadjuvanted vaccine (GMT 1:22,267,288 and 1:129,140,163 vs 1:27 respectively; **Figure 1**). Mice vaccinated with gp120+M7 had significantly elevated serum anti-gp120 IgA titers (GMT 1:2,187; 4 of 5 mice developed serum anti-gp120 IgA) while only 2/5 mice vaccinated with gp120+CT and 1 of 5 mice vaccinated with gp120+MPL developed serum anti-gp120 IgA titers (**Table 3**). Serum anti-gp120 IgA was not detected in mice that received gp120 alone or gp120+c48/80. Additionally, only mice that received vaccines adjuvanted with c48/80 or M7 developed significantly elevated vaginal gp120-specific IgG responses (1:1176 and 1:588, respectively), Only mice vaccinated with gp120+M7 developed significantly elevated vaginal gp120-specific IgA titers (GMT 1:73; 4 of 5 mice developed vaginal anti-gp120 IgA., (**Table 3**). Based on these results, M7 was selected as the nasal vaccine adjuvant for subsequent experiments in rabbits and NHPs.

**Figure 1.**
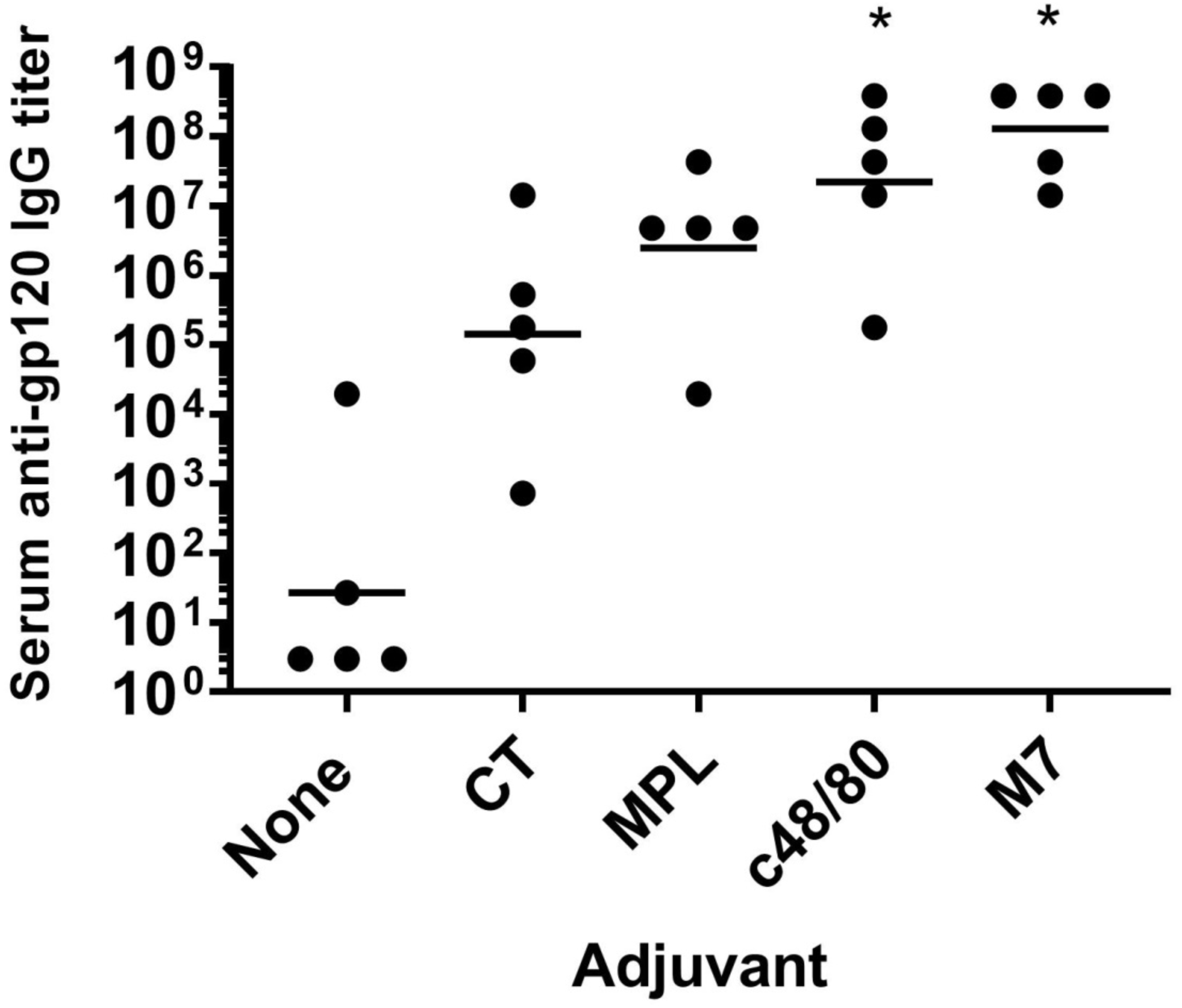
Intranasal vaccines adjuvanted with the mast cell activating adjuvants compound 48/80 or Mastoparan 7 induce stronger serum gp120-IgG titers than induced by cholera toxin or monophospholipid. **A.** C57BL/6 mice (n=5/group) were vaccinated on day 0, 7 & 21 with 10µg gp120 and either no adjuvant, 5nM M7, 15µg c48/80, 1µg CT, or 10µg MPL. Serum was collected on day 35 and gp120-specific IgG titers determined by ELISA. Mice receiving vaccines adjuvanted with c48/80 or M7 had significantly higher serum anti-gp120IgG titers than mice vaccinated with gp120 alone (p=0.01, 0.001 respectively). The horizontal line indicates the geometric mean titer.

### Intranasal immunogenicity of HIV-1 gp120 and gp120-Ad2F

One goal of this study was to evaluate mucosal immunization for its ability to induce serum anti-HIV-1 antibody titers similar to those induced by IM immunization. For mucosal immunization with HIV-1 gp120, IN immunization was utilized due to our previous success using this route in non-human primates (8, 60). Since the rabbit nasopharyngeal lymphoid tissues are similar with those in primates and humans (90-92), results from the rabbit model may be more likely to translate to NHPs and humans than results obtained from mice. Thus, rabbits were used as the animal model to evaluate intranasal immunization with HIV-1 gp120 combined with the cationic antimicrobial mast cell activating peptide M7.

Higher antigen doses are likely required for intranasal immunization to induce circulating antibody responses similar to those induced by injected vaccines (9). Therefore, we tested IN vaccination with 100, 200, and 300 µg of gp120 adjuvanted with the M7 peptide to determine what dose of gp120 administered IN was required to achieve serum anti-gp120 IgG titers similar to those induced by IM vaccination with 100 µg of HIV-1 gp120 adjuvanted with Addavax, a squalene-based oil-in-water emulsion (see experiments IN 1-3 in **Table 1**). Squalene oil-in-water adjuvanted vaccines induce titers similar to or higher than alum adjuvanted vaccines (61, 62) and our previous success using MF59 (a squalene-based adjuvant) in a prime-boost vaccine regimen led to our use of Addavax in the current study (8).

As shown in **Figure 2**, while only 4 of 8 animals that received 100 µg of gp120 IN developed detectable serum anti-gp120 IgG titers on day 42 after two vaccines, all rabbits in the remaining groups at this time point exhibited detectable serum gp120-specific IgG. Rabbits that received 100 µg or 200 µg gp120 IN developed serum anti-gp120 IgG titers that were significantly different from 100 µg IM (p<0.0001, p=0.0015 respectively). Thus, we considered 300 µg gp120 administered IN to induce serum gp120-specific IgG titers similar to 100 µg gp120 administered IM.

**Figure 2.**
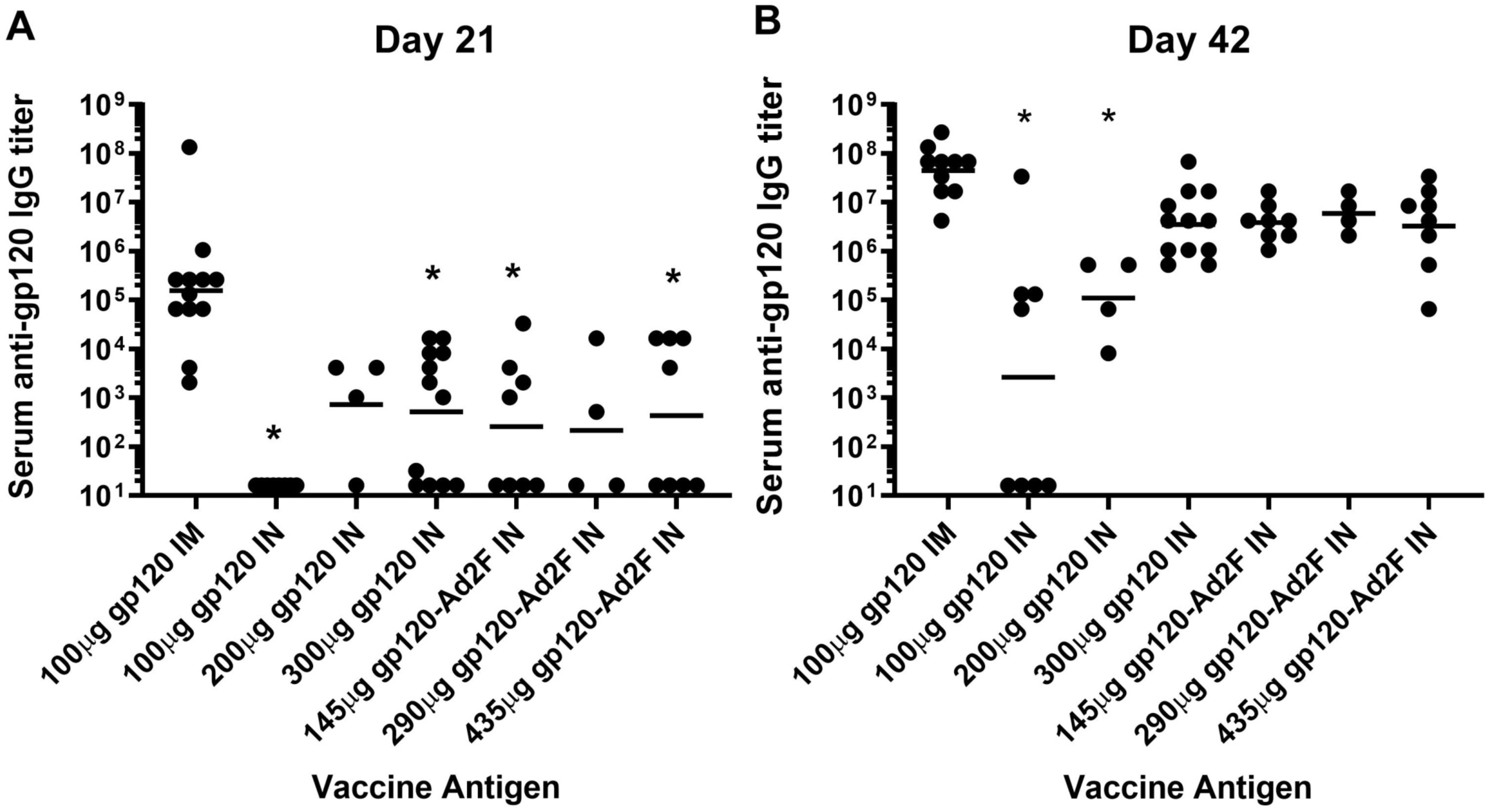
300µg gp120 and gp120-Ad2F (equimolar doses to 100-300µg gp120) administered IN induce serum anti-gp120 IgG titers similar to intramuscular administration of 100µg of gp120. Male New Zealand White rabbits were vaccinated intramuscularly with 100µg gp120 adjuvanted with Addavax or intranasally with gp120 or gp120-Ad2F adjuvanted with 64µg M7. The intranasal vaccines contained either 100, 200, or 300µg gp120 or 145, 290, or 435µg of gp120-Ad2F (doses equimolar to gp120). Serum was collected two weeks after the first vaccination (day 21; A) and the second vaccination (day 42; B) and assessed for gp120-specific IgG responses by ELISA. Each group has an n=4 and was repeated 0-2 times for a total n of 4-12. A) After a single vaccination, serum anti-gp120 IgG responses induced by 100 and 300ug of gp120 or equimolar doses of Ad2F were significantly lower than anti-gp120 IgG responses induced by 100µg gp120 IM (p<0.05). B) After administration of two vaccines, only 100µg and 200µg gp120 administered intranasally induced serum anti-gp120 IgG responses significantly lower than serum ant-gp120 IgG responses induced in the 100µg gp120 IM group. (p<0.0001 and 0.001 respectively). The horizontal line indicates the geometric mean titer.

We also tested gp120-Ad2F at doses (145, 290, and 435µg) that were equimolar to the IN doses of gp120 tested, to determine if the addition of the mucosal targeting ligand Ad2F to gp120 enhanced the immunogenicity of gp120 when IN delivered (experiments IN 2 & 3 in **Table 1**). In contrast to the results obtained with gp120, none of the gp120-Ad2F vaccine groups induced serum anti-gp120 IgG titers that were significantly different from the anti-gp120 IgG titers induced by the gp120 IM vaccine group after two immunizations (**Figure 2B**). However, the geometric mean gp120-specific IgG titer in all of the gp120-Ad2F vaccine groups was a log lower than the IM group. Additionally, regardless of gp120-Ad2F dose, only 50% of rabbits in each group had a detectable anti-gp120 IgG antibody response after the first dose (4/8, 2/4, 4/8) while all 12 rabbits that received 100 µg gp120 IM had a detectable antibody response after the first vaccination (**Figure 2A**). A dose response was not observed for gp120-Ad2F vaccine groups at either time point. Since gp120-Ad2F was more immunogenic than gp120 by more than two orders of magnitude at the 100µg dose, gp120-Ad2F was used as the antigen for the remaining intranasal immunization studies.

### Evaluation of the mucosal immunogenicity of MVA expressing gp120

We tested various forms of MVA vectors expressing gp120 to determine if vector modifications enhanced their mucosal immunogenicity as determined by serum anti-gp120 IgG responses induced. The immunogenicity of the replication-defective MVAdelta5 vector expressing gp120 (MVAdelta5gp120), similar to thatdeveloped by Garber et al. (42), was first compared to the immunogenicity of conventional MVA expressing gp120 (MVAgp120) when delivered by the IM route (MVA 2 and MVA 3 in **Table 1**). A dose response for both anti-gp120 IgG and vector-specific IgG (anti-B5, anti-L1) was observed with escalating doses of MVAdelta5gp120 (**Figure 3**). 1×10^10^ PFU of MVAdelta5gp120 induced anti-gp120 IgG titers most similar to anti-gp120 IgG titers induced by immunization with 1×10^8^ PFU of MVAgp120. However, MVAgp120 induced a more consistent gp120-specific IgG response than MVAdelta5gp120, potentially indicating an even higher dose of MVAdelta5gp120 may be needed to induce consistently high anti-gp120-specific titers in all individuals.

**Figure 3.**
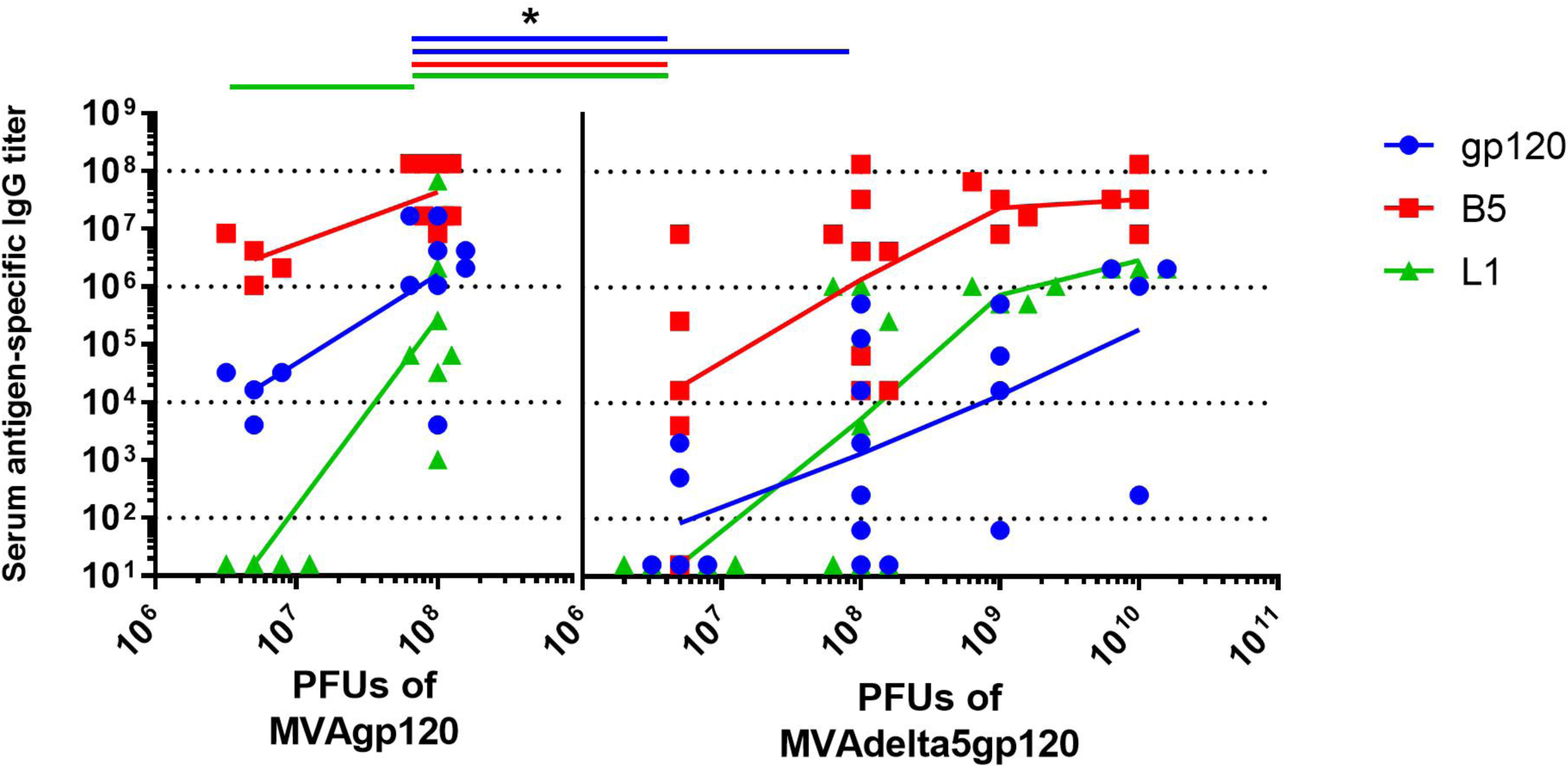
Non-replicating MVAdelta5gp120 is not superior to conventional MVAgp120 for induction of serum gp120-specific IgG. Female rabbits were intramuscularly vaccinated with MVAgp120 or MVAdelta5gp120 at the doses indicated on day 0 & 21. Rabbits were bled on day 35 and gp120-specific and vector specific (B5 and L1) IgG was determined by ELISA. * Significantly different from 1×10^8^PFU of conventional MVA (p<0.05). The connecting lines represent the geometric mean titer.

After determining that MVAgp120 was the most immunogenic form of MVA to use as a priming vaccine, different mucosal routes of immunization with MVAgp120 were evaluated for their ability to induce mucosal and systemic anti-HIV-1 gp120 antibody responses. Based on needing three times the protein dose for gp120 IN vaccines to induce similar serum antibodies as IM vaccination, a dose of 3×10^8^ PFU of MVAgp120 administered by various mucosal routes (nasal, rectal, and gastric) was used for comparison to 1×10^8^ PFU IM (MVA 1 in **Table 1**). After the second vaccination, all IM vaccinated rabbits had detectable gp120-specific IgG titers while only 2/3 of IN vaccinated rabbits had detectable gp120-specific IgG titers (**Figure 4**). No rabbit that was rectally or gastrically immunized developed detectable serum gp120-specific IgG titers (<1:32 titer). Based on these data, IN prime with MVAgp120 was selected for use in the prime/boost experiments.

**Figure 4.**
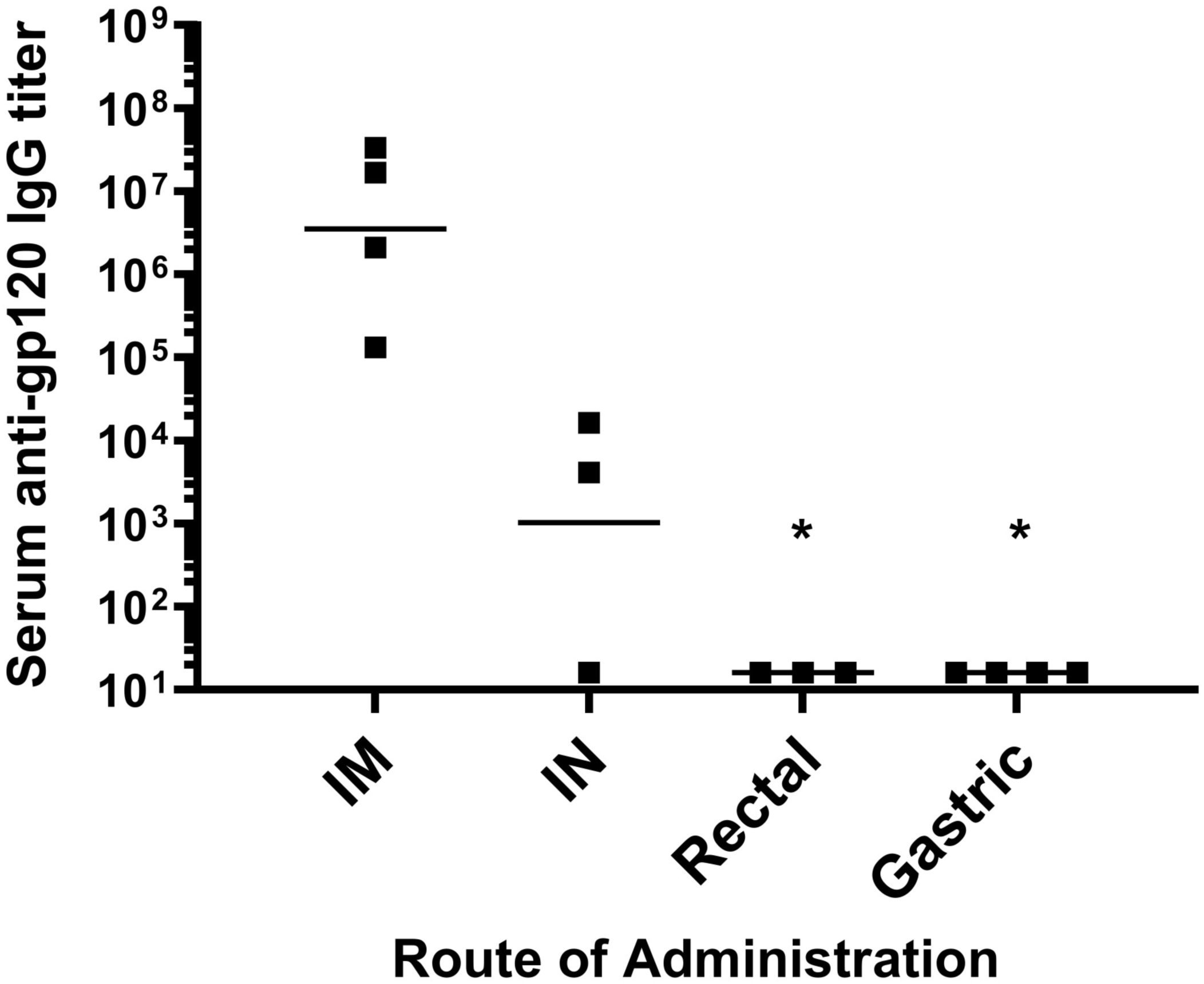
Intranasal but not rectal or gastric administration of MVAgp120 induced serum gp120-specific IgG titers. Female New Zealand White rabbits were vaccinated with MVAgp120 by the intramuscular, intranasal, intragastric, or intrarectal routes (n=3-4/group) on days 0 and 28. Serum was collected on day 42 and gp120-specific IgG titers were determined by ELISA. Rectal and gastric administration of MVAgp120 resulted in serum gp120-specific titers that were significantly lower than IM administration (p=0.01, 0.008 respectively). The horizontal line indicates the geometric mean titer.

### Evaluation of the immunogenicity of MVA-gp120 prime / gp120 boost immunization regimens in rabbits

Three different prime/boost regimens that included nasal vaccination were compared to an intramuscular prime/boost regimen to determine if a prime/boost regimen utilizing IN immunization could be developed that induced serum anti-gp120 IgG responses similar to those induced by the IM prime/boost regimen We also evaluated the use of a boosting regimen that utilized a combination of both the IM and IN routes since combining the two routes of administration enhanced mucosal antibodies and enhanced serum viral neutralization activity in previously vaccinated NHPs (46). The prime/boost regimens evaluated that included nasal immunization include IM MVAgp120 prime, IN gp120-Ad2F boost (IM/IN); IM MVAgp120 +prime, IN gp120-Ad2F + IM gp120 boost (IM/IM+IN) and IN MVAgp120 prime with IN gp120-Ad2F + IM gp120 boost (IN/IM+IN; see **Table 2**). At the end of the prime/boost regimen (week 19), only the IM/IN group had significantly lower (p=0.01) serum gp120-specific IgG titers than the control IM/IM group (**Figure 5A**). There were no significant differences in serum V1V2-specific IgG titers (**Figure 5B**), serum anti-gp120 IgA titers, or gp120-specific fecal or vaginal antibodies (**Figure 6**). Additionally, there were no significant differences in the gp120-specific ADCC titers between groups (p=0.19). However, the geometric mean ADCC titer was most similar between the IM/IM (1:24,705) and the IN/IM+IN group (1:33,921) than the IM/IN (1:8,591) or IM/IM+IN (1:48,626) groups (**Figure 5C**). However, the IM/IM+IN vaccinated rabbits developed significantly higher neutralization of 6644.v2.c33 (p=0.04), a tier 1b pseudovirus, than the IM/IM group (**Figure 5D**). The IN/IM+IN vaccination regimen was employed for further comparison to the IM/IM regimen in NHPs as these groups had the most similar functional serum antibody responses in rabbits and allowed a comparison of IM prime/boost to a vaccine regimen that utilized IN priming and boosting to maximize the use of mucosal immunization.

**Figure 5.**
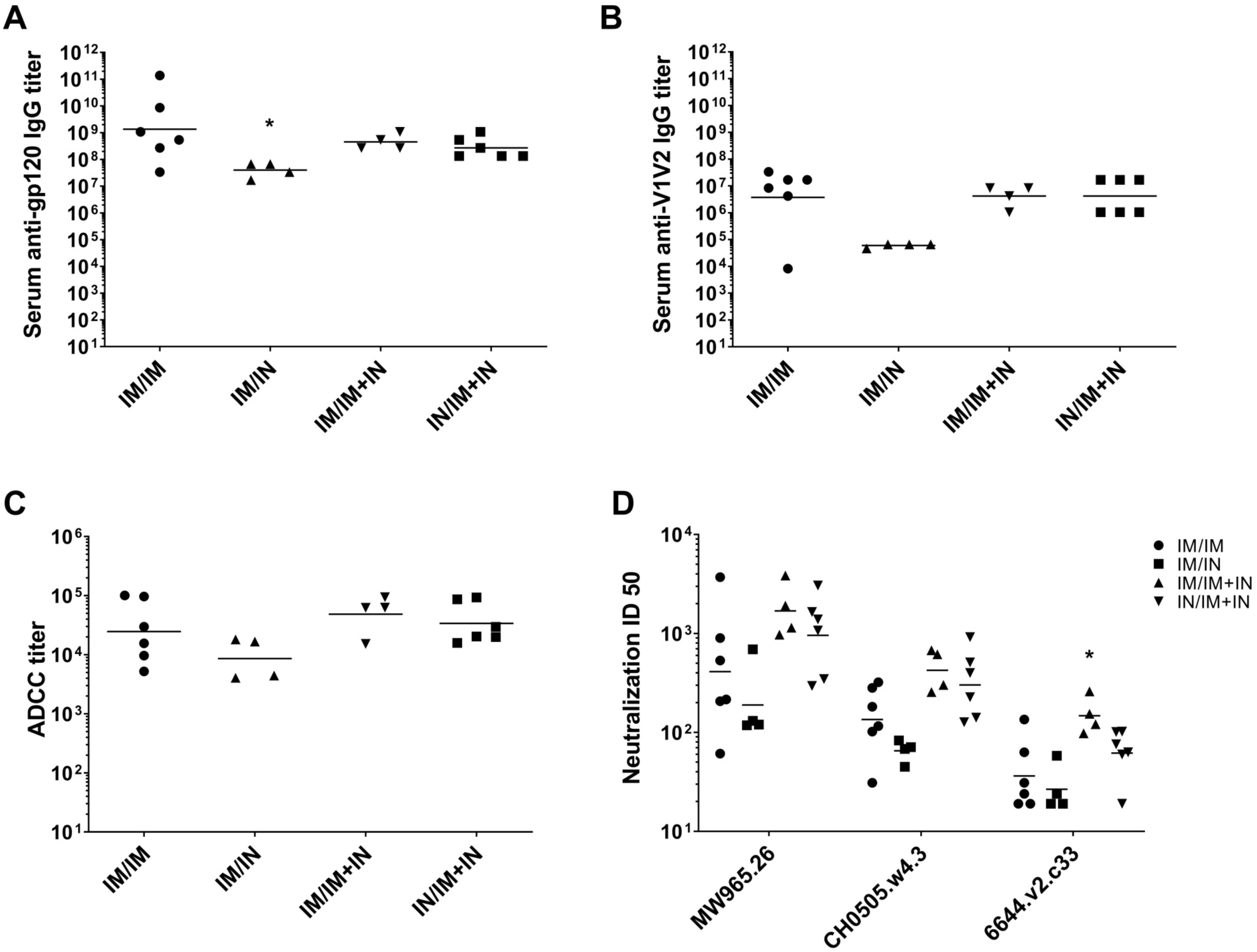
Combined intramuscular and intranasal boosting, regardless of an intramuscular or intranasal prime, induces serum anti-HIV antibodies similar to those induced byintramuscular prime/boost regimen in rabbits. New Zealand White rabbits were primed intramuscularly(IM) or intranasally (IN) with MVAgp120 on week 0 and boosted with 100µg gp120 adjuvanted with Addavax IM and/or with 435µg gp120-Ad2F adjuvanted with 64µg M7 IN on weeks 12 and 16. N=4-6/group. Serum antibody titers and viral neutralization from blood drawn on week 19 shown. A) Serum gp120-specific IgG titers. Serum anti-gp120 IgG titers induced by IM/IN is significantly lower than serum anti-gp120 IgG titers induced by IM/IM (p=0.01). B) Serum V1V2-specific IgG titers were not significantly different between groups. Kruskal-Wallis was not significant (p=0.057) so multiple comparisons were not made. C) ADCC activity using human effector cells against HIV gp120-coated target cells. No significant differences between groups (p=0.19). D) Neutralization of tier 1a (MW965.26 and CH0505.w4.3) and tier 1b (6644.v2.c33) pseudoviruses. IM/IM+IN immunization induced significantly higher neutralization of 6644.v2.c33 (p=0.04) than immunization by the IM/IM route. The horizontal line indicates the geometric mean titer.

**Figure 6.**
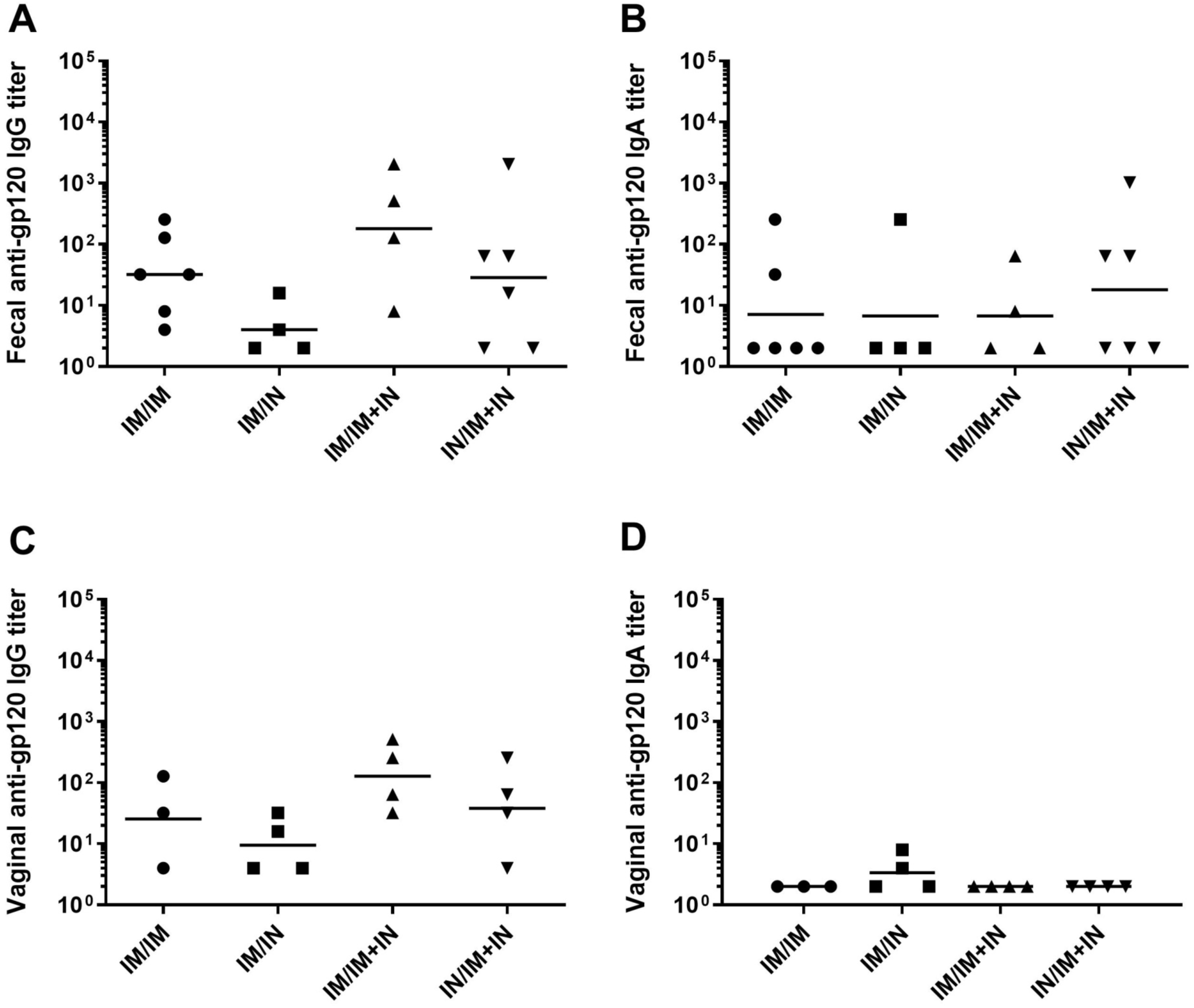
Rabbits developed undetectable or low mucosal antibody responses to all of the prime/boost regimens. New Zealand White rabbits received a priming immunization of MVAgp120 either IM or IN on week 0 and gp120 boosts IM and/or IN on weeks 12 and 16. Feces were collected from all individuals (n=4-6/group) while vaginal lavage was only performed on the females (n=3-4/group). Results shown are from week 19, three weeks after administration of the second booster vaccine. A) 50-100% of rabbits in each group developed detectable fecal gp120-specific IgG titers. B) 25-50% of rabbits in each group developed fecal gp120-specific IgA titers. C) Vaginal gp120-specific IgG was detectable in 50-100% of the female rabbits in each group but tended to be lower than the fecal gp120-specific IgG titers observed. D) Vaginal gp120-specific IgA was only detectable in 2 of the 4 IM/IN vaccinated females. The horizontal line indicates the geometric mean titer.

### Evaluation of the immunogenicity of the IN/IM+IN prime/boost vaccine regimen in NHPs

To determine if a combined mucosal/systemic vaccine regimen optimized in rabbits is translatable, NHPs were vaccinated with the control IM/IM regimen (n=6) or with IN/IM+IN regimen (n=6) developed in rabbits to determine if the IN/IM+IN regimen was able to induce serum anti-gp120 IgG responses similar as the IM/IM regimen (see **Table 2**). The vaccination regimens induced antibody responses in the NHPs that were similar to the antibody responses induced by the same vaccine regimens in rabbits (**Figure 7**). There was no significant difference in week 19 serum gp120-specific IgG or IgA, serum V1V2-specific IgG, fecal gp120-specific IgG or IgA titers, or in virus neutralization between IM/IM and IN/IM+IN regimens in the NHPs (**Figure 7&8**). As reported by Pollara et al (accepted JVI, JVI02119-18R1, Bridging vaccine-induced HIV-1 neutralizing and effector antibody responses in rabbit and rhesus macaque animal models), there were no significant differences in ADCC activity against gp120-coated target cells or antibody dependent cellular phagocytosis (ADCP) of HIV-1 virions between the two vaccination regimens.

**Figure 7.**
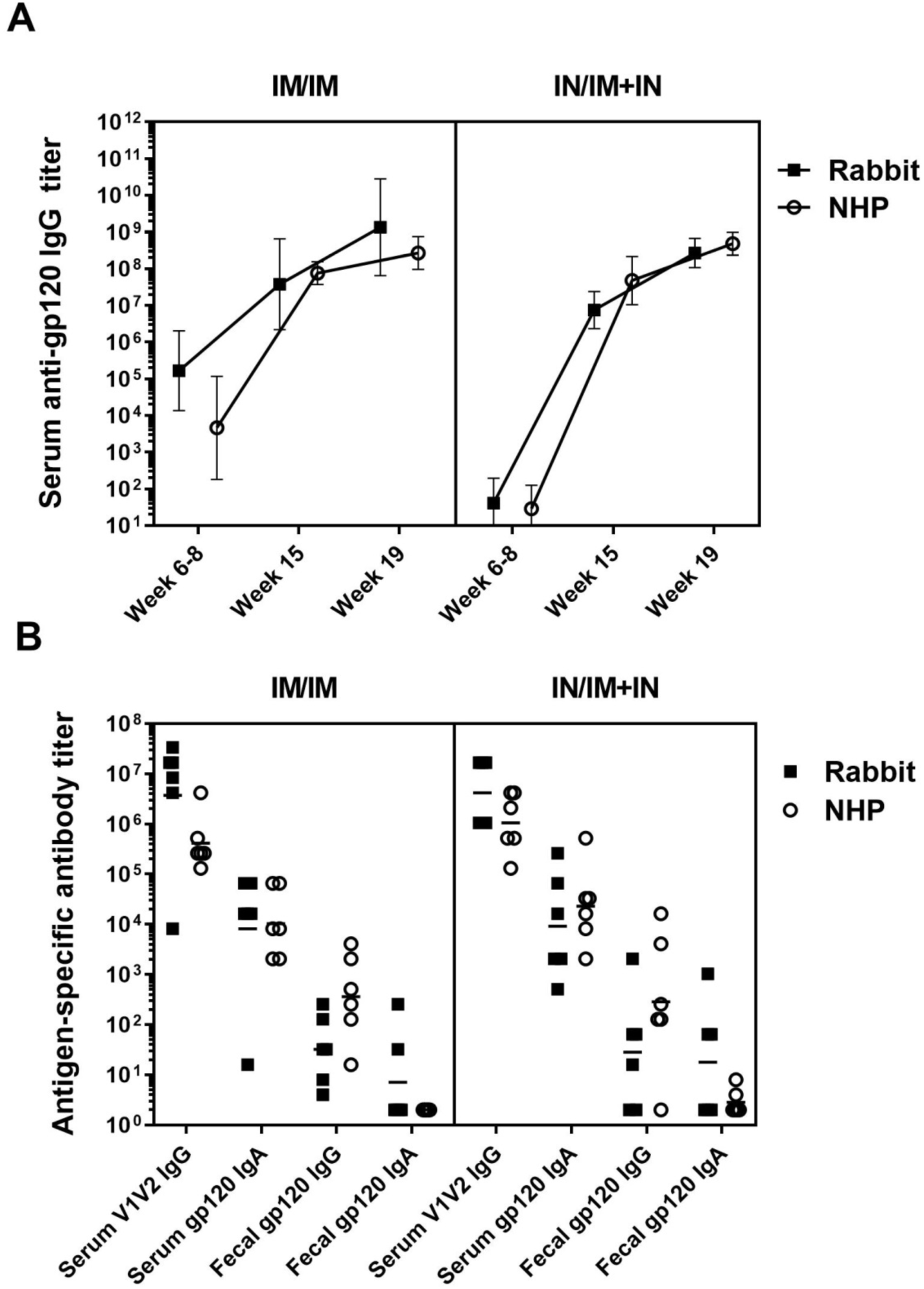
Rabbit and NHP antibody responses were similar to both prime/boost regimens. New Zealand White rabbits and rhesus macaques received a priming immunization of MVAgp120 on week 0 and gp120 boosts on weeks 12 and 16. All vaccines were either administered intramuscularly (IM prime/IM boost) or the MVAgp120 prime was administered intranasally and boosting consisted of both an intramuscular and intranasal component (IN/IM+IN)(n=6/group). Animals were bled at the timepoint indicated and antibody responses were determined by ELISA. A) Serum gp120-specific IgG responses after administration of the MVAgp120 prime (weeks 6-8), the first gp120 boost (week 15), and the second gp120 boost (week 19). The connecting lines and error bars represent the geometric mean titer and the 95% confidence intervals. B) After administration of the second booster vaccine (week 19) rabbits and NHPs had similar serum V1V2-specific IgG and serum gp120-specific IgA responses. Fecal gp120-specific responses were also similar between species. The horizontal lines represent the geometric mean titer.

NHPs received a third booster vaccine corresponding with their assigned vaccination regimen (thus IM or IM+IN) on week 24. Prior to administration of the 3^rd^ booster, serum gp120-specific IgG titers were significantly greater (p=0.03) in the IN/IM+IN group compared to the IM/IM group, potentially indicating that IN/IM+IN serum anti-gp120 IgG responses are more durable that IM/IM induced responses. However, there was no significant difference in serum gp120-specific IgG or IgA titers after the 3^rd^ boost (weeks 27 and 30) (**Figure 8a,b**). There was also no significant change in fecal anti-gp120 IgG titers after the third boost (**Figure 8c**). Fecal gp120-specific IgA was undetectable in any week 19 sample at a 1:16 dilution, but was detected in week 27 samples at dilutions ≤ 1:32 for 2/6 primates in each vaccine group (**Figure 8d**). Our results indicate that a prime/boost vaccine regimen utilizing IN immunization can be developed that induces anti-HIV-1 serum IgG and IgA and fecal IgG responses similar to those induced by IM/IM immunization.

**Figure 8.**
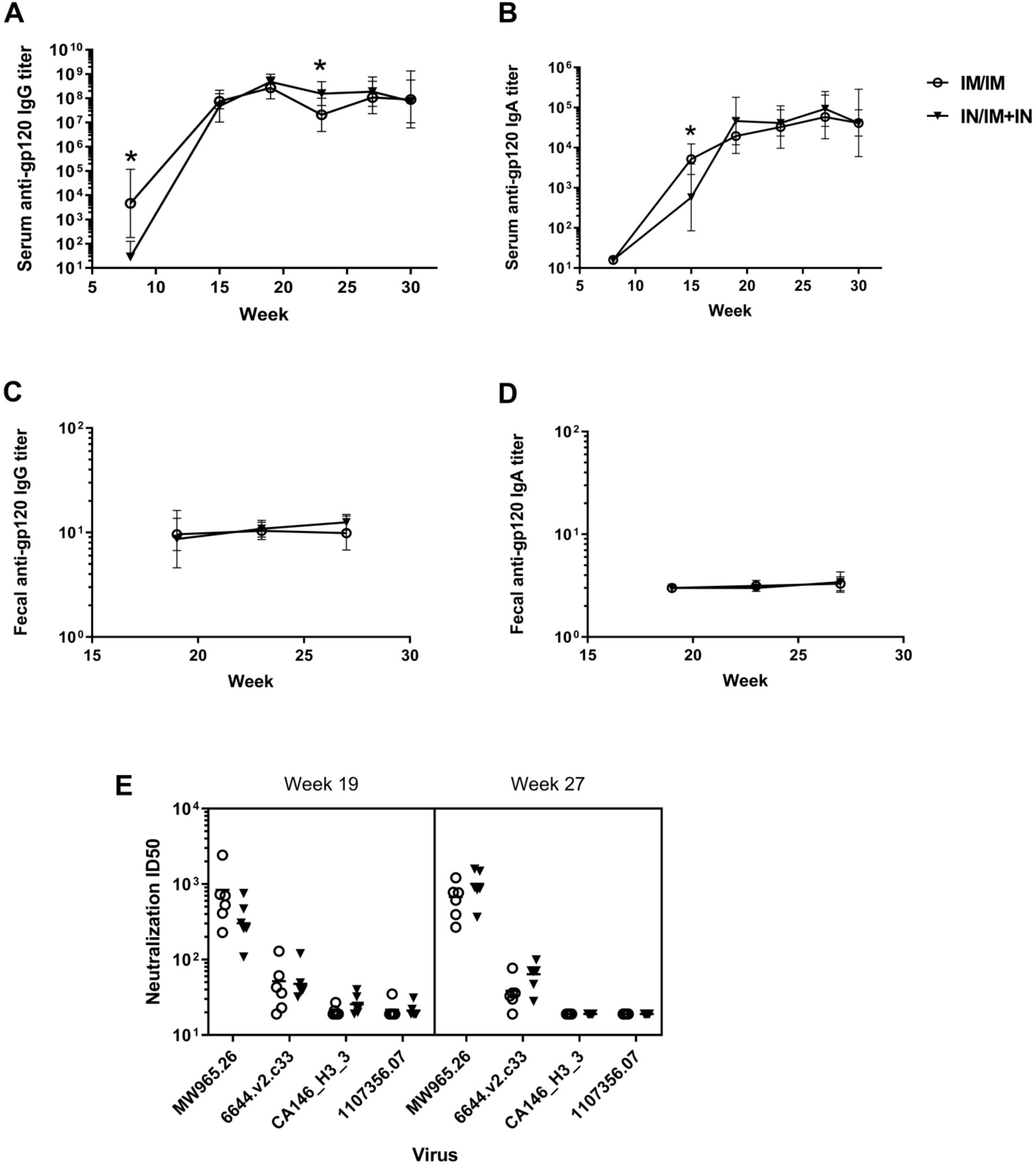
Intranasal prime with combined intranasal and intramuscular boosting induces similar serum antibodies and viral neutralization as the parenteral IM prime/IM boost regimen. Rhesus macaques (n=6/group) were vaccinated with the same IN/IM+IN or IM/IM regimens as described for rabbits. An additional booster vaccine was given on week 24. Serum was collected at indicated timepoints and antigen-specific antibody titers were determined by ELISA. Antibody responses were compared between the two vaccination regimens at each timepoint using a Mann-Whitney test. A) Time course of serum gp120-specific IgG titers during the vaccination regimen. IM/IM had significantly higher serum gp120-specific IgG after administration of the priming immunization (week 8) while IN/IM+IN had significantly higher titers 7 weeks after administration of the second booster vaccination (week 27) (p=0.01, 0.03 respectively). The connecting lines and error bars represent the geometric mean titer and the 95% confidence intervals. B) Time course of serum gp120-specific IgA titers. The IM/IM regimen resulted in significantly higher anti-gp120 IgA titers than the IN/IM+IN regimen at week 15 (p=0.04). The connecting lines and error bars represent the geometric mean titer and the 95% confidence intervals. C) Fecal gp120-specific IgG was not significantly different between vaccination regimens at any time point tested. The connecting lines and error bars represent the geometric mean titer and the 95% confidence intervals. D) While all NHPs had fecal gp120-specific IgA undetectable at a 1:16 dilution at week 19, low levels of fecal gp120-specific IgA were detected in a few NHPs in each vaccination regimen on week 27. The difference between timepoints was not significant (p=0.6 for IM/IM, p=0.3 for IN/IM+IN). The connecting lines and error bars represent the geometric mean titer and the 95% confidence intervals. E) Neutralization of tier 1a (MW965.26) and tier 1b (all others) pseudoviruses was not significantly different between the vaccination regimens after both the 2^nd^ and the 3^rd^ boost. The horizontal lines represent the geometric mean titer.

### Evaluation of protective immunity using a repeated low-dose rectal SHIV challenge

The NHP vaccinated using the IM/IM and IN/IM+IN regimens received a repeated escalating dose rectal challenge with a heterologous tier-2 SHIV on week 30 to determine if the use of mucosal immunization routes provided any protective benefit vs the IM/IM vaccine regimen. Four unvaccinated NHPs were challenged as negative controls. Following the four low dose challenges, challenge doses were increased by an order of magnitude every other week to determine if the vaccination regimens resulted in protection. All unvaccinated animals became infected by the 7^th^ +/-1 challenge (**Figure 9A**). 2 of the 6 NHPs in both the IM/IM and IN/IM+IN groups became infected during the initial low dose challenges. All of the IM/IM group and 5/6 individuals in the IN/IM+IN group became infected on or by the 7^th^ challenge. Thus, there was no significant difference in number of challenges until infection between vaccine groups. Additionally, similar viral loads were present in the immunized and unimmunized animals with peak viral load two weeks post-infection (**Figure 9B**). The lack of protection seen in both the IM/IM and IN/IM+IN vaccine prevents us from determining if the use of an optimized mucosal vaccine regimen provides superior protection against a mucosal SHIV challenge.

**Figure 9.**
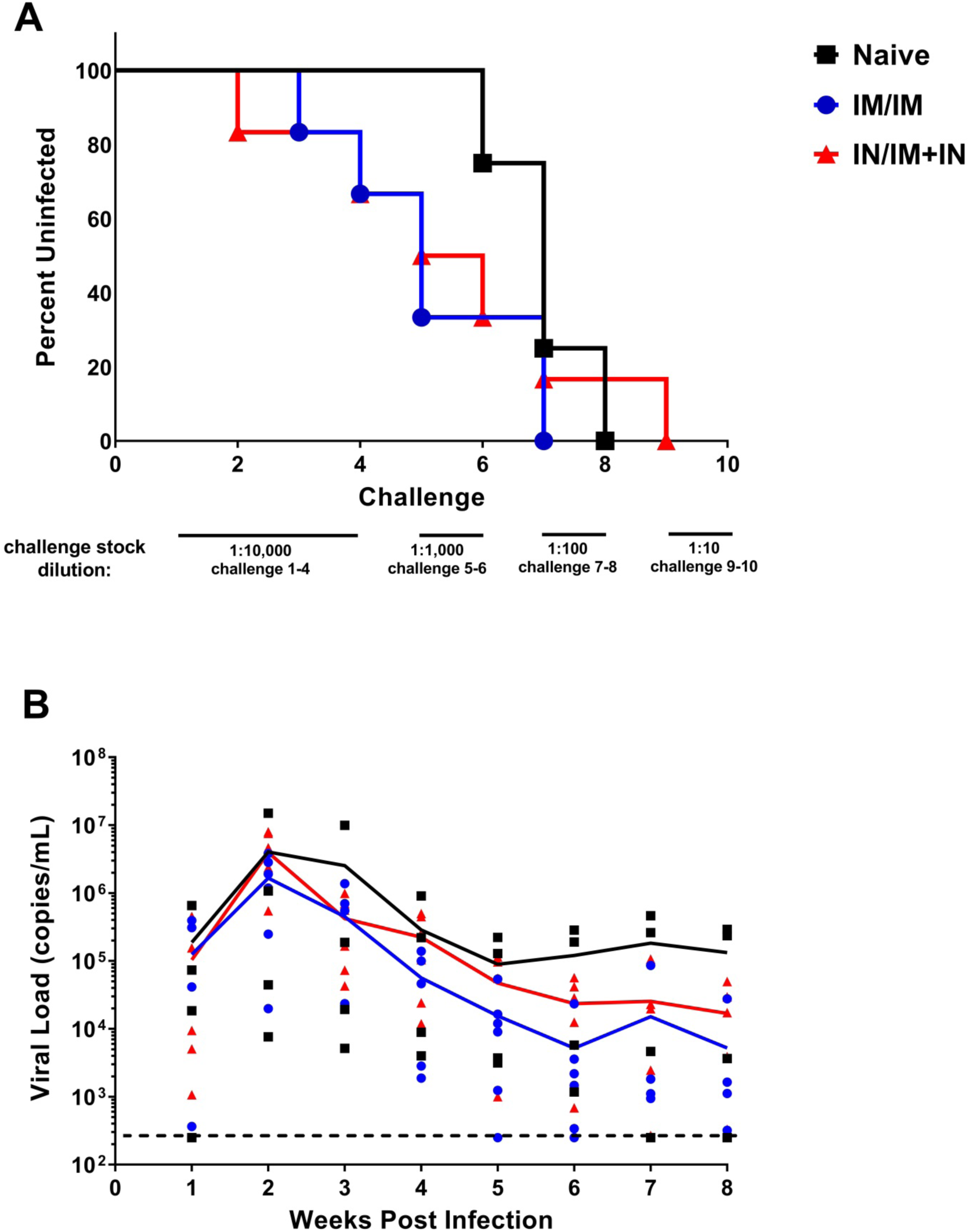
Neither the IM/IM vaccine regimen nor the optimized IN/IM+IN regimen protected NHPs from SHIV challenge. Rhesus macaques (n=6/vaccine group; n=4/naïve group) were rectally challenged weekly with an escalating dose of heterologous tier-2 SHIV-1157QNE Y173H. All animals received two additional challenges after the infective challenge or a maximum of 10 challenges. Blood was collected one week after challenge (same day as administration of next challenge) and assessed for viral RNA by PCR. A) SHIV infection status after administration of each challenge. After five challenges, 4 of 6 individuals that received the IM/IM regimen and 3 of 6 individuals that received the IN/IM+IN regimen were infected while 0 of 4 unvaccinated individuals were infected. There was no statistical difference between vaccinated and naïve animals (p=0.08, Fisher’s exact test) even at that time point. B) Viral load after infection as determined by PCR. Line represents the mean viral load for the vaccine group. Data points represent individual’s viral load for that timepoint. Limit of detection is 250 copies/mL.

## DISCUSSION

The use of mucosal immunization may induce antigen-specific mucosal IgA responses that are significantly greater than antigen-specific mucosal IgA responses induced by parenteral immunization(8), while the serum antigen-specific IgG binding antibodies(9) and functional antibodies(8) induced by mucosal immunization are often lower than those induced by parenteral immunization. Since protection against HIV-1 mucosal transmission may require optimum systemic and mucosal anti-HIV-1 antibody responses(7), we propose that HIV-1 vaccine regimens utilizing a mucosal route of immunization should be optimized to ensure induction of elevated serum anti-HIV-1 binding and functional antibody responses. In this study we demonstrate that a prime-boost HIV-1 immunization regimen utilizing IN immunization may be optimized to induce serum anti-HIV-1 binding and functional antibodies at a quantity similar to that induced by prime-boost immunization regimen delivered by the IM route.

Here we report for the first time the use of M7, a cationic antimicrobial and mast cell activating peptide, as an adjuvant in a HIV vaccine formulation. M7 provided more potent adjuvant activity than CT, the “gold-standard” for nasal vaccines. The efficacy of M7 is not surprising as vaccines adjuvanted with compound 48/80, another mast cell activator, have induced antibody titers in mice and rabbits similar to CT-adjuvanted vaccines (14, 17, 63). Others have demonstrated that a vaccine formulation including an antimicrobial peptide adjuvant, IC31, was an effective adjuvant for the induction of potent anti-HIV-1 immune responses in mice after parenteral immunization (64). Our results suggest that M7 is an effective nasal vaccine adjuvant that could be evaluated in future human IN HIV-1 vaccine studies althoughadditional safety studies are needed with M7 prior to its use in clinical studies.

It is documented in the mucosal vaccine field that higher antigen doses are needed for mucosal vaccines to induce antibody responses similar to those induced by injected vaccines (65-67). Consistent with this, we demonstrated that IN administration of 300µg of gp120 was required to induce serum gp120-specific IgG titers not significantly different from those induced by IM administration of 100µg of gp120. Others have reported the use of 3-fold higher antigen doses for IN vaccination did not induce serum antibody responses similar to those induced by parenteral immunization. For example, a NHP study that administered five immunizations with either 300µg of gp140 adjuvanted with LTK63 administered IN or 100µg of gp140 adjuvanted with MF59 administered IM, failed to achieve similar serum gp140-specific IgG titers between the two groups (68). Other studies have compared IN to systemic vaccination using the same antigen dose (9, 69, 70), two-fold higher (71), three-fold higher (68), or five-fold higher (9). Differences in the immunogenicity of different nasal vaccine regimens may be attributable to antigen (specific antigen and dose), adjuvant (specific adjuvant and dose), and the method of nasal vaccine delivery(72) and the vaccine volume(73) administered to a nostril at a given time.

It is important to note that we did not determine that 100µg gp120 IM and 300µg gp120 IN are equivalent, but rather we failed to detect a significant difference. Increasing statistical power by increasing the sample size and meeting the assumptions for parametric statistics would increase the likelihood of detecting a difference between these two vaccines. We consider the variation in individual responses observed with the use of an outbred rabbit model beneficial for translation to real world use where vaccines are administered by a large number of individuals to people with different immune status, genetic backgrounds, and environmental exposures. For example, the extremely large variation in gp120-specific titers after vaccination with 100µg of gp120 is consistent with a response along the linear portion of a dose response curve indicating that slight differences in administration or in immune responses between animals could determine if the vaccine induced an immune response or if the animal was a non-responder. Notably, the 100µg gp120 group was repeated, with 4 rabbits per replicate and different personnel administering the IN vaccines in each replicate, and there were responders and non-responders in each replicate. Since statistical power and sources of variance are important and often overlooked factors when evaluating comparisons of nasal and systemic vaccines, our results may perhaps more accurately be viewed as demonstrating a difference between 100µg gp120 IM and 100µg or 200µg gp120 IN.

The inclusion of the cell adhesin, Ad2F, greatly improved the immunogenicity of gp120. IN vaccination with 145µg of gp120-Ad2F, which is equimolar to 100µg gp120, induced serum antibody titers similar to the 100µg gp120 IM administered while IN vaccination with 100 µg gp120 induced much lower and more variable anti-HIV-1 IgG responses. The approximately 100-to 1,000-fold increase in serum gp120-IgG titers induced by using gp120-Ad2F vs. gp120 is similar to that observed with other Ad2F-coupled immunogens delivered by the nasal route in rabbits(17). While we did not see a dose response effect when increasing to doses equimolar to 200µg gp120 or 300µg gp120, we hypothesize that this is due to reaching the upper plateau of the dose response curve. If this is indeed the case, it is possible that even doses lower than 145µg of gp120-Ad2F could be used and still achieve high serum antibody titers, allowing for antigen sparing. Conversely, there may be a benefit to using the gp120-Ad2F dose equimolar to 300µg gp120, for example reduced variation in titers, that this study was not sufficiently powered to detect. Since recombinant adenovirus vectors have been safely delivered by the nasal route to humans (74, 75), it is hopeful that immunogens containing Ad2F as a mucosal targeting ligand will be safe for human use.

To optimize the MVA vectored priming immunization, the effects of modifications to the MVA vectors through the use of a replication-defective MVA containing deletions of four genes encoding immune-modulatory proteins was evaluated. Although others have reported the induction of similar vector-specific and insert-specific antibody titers with replication-defective and conventional MVA vectors (39), in this study 1×10^8^ PFUs of MVAdelta5gp120 induced significantly lower serum gp120-specific IgG responses than an equivalent dose of the conventional MVAgp120 vector. The lower response may be explained by differences between studies including: animal models (NHPs vs rabbits; route of administration (ID+IM vs IM), HIV antigen (gag vs gp120); how ELISA titers were determined (OD of single serum dilution vs endpoint titer determined from serial dilutions); and most notably, differences in the promoters used to drive expression of HIV antigens. In the previous studies the modified H5 promoter was used to provide early transcription and reduced late transcription (42), whereas in this study, the gp120 expression was controlled by a synthetic promoter designed to promote strong late gene expression as well as early gene expression. Thus, in this present study, the replication-defective MVA vector would be expected to express less gp120 protein than the conventional MVA, because in the absence of the viral uracil-DNA-glycosylase and DNA replication, late expression would be abrogated.

The intranasal route was identified as the most effective mucosal route for priming with MVA, as determined by serum anti-gp120 IgG similar responses as the intramuscular regimen after administration of the first protein booster vaccine, when combined IN/IM gp120 boosts were administered. This result is similar to what was observed with an HIV peptide vaccine, as in both studies nasal vaccination was the most immunogenic mucosal route (49). Even though IN administration of MVA was effective for priming, similar immunogenicity between the IM and IN priming immunization was not achieved, even when three times more MVA was IN administered. Regardless, there was no significant difference in serum antibody titers after administration of the first boost indicating that IN administration of MVA was an effective priming immunization. Our results are consistent with a previous study in NHPs where there was a significant difference in serum IgG titers when MVA was administered IM versus IN, however, nasal vaccination resulted in higher T cell responses and a longer interval until progression to AIDS than the IM vaccination (76).

Using the MVA prime (IM or IN) and IN gp120-Ad2F and IM gp120 vaccine delivery, we tested various combinations of systemic and IN prime and boost vaccines including IM MVA prime + IM gp120 boost (IM/IM),, IM MVA prime + IN gp120-Ad2F boost (IM/IN), IM MVA prime + IM gp120 and IN gp120-Ad2F combination boost (IM/IM+IN), and IN MVA prime + IM gp120 and IN gp120-Ad2F combination boost (IN/IM+IN). In our rabbit model, only the IM/IN regimen resulted in significantly lower serum gp120-specific IgG titers than the IM/IM regimen. A previous NHP study did not observe a significant difference in plasma anti-gp120 IgG binding responses between a similar vaccination regimen consisting of an MVA prime and two gp120 boosts administered either IM or IN(8). However, differences in plasma virus neutralization between the IM/IM and the IM/IN vaccinees indicates that the IM/IN regimen induced lower functional IgG responses than the IM/IM regimen, congruent with our results. Additionally, NHPs from the IM/IN regimen in the previous study went on to receive a combined IM+IN boost almost a year after the MVA prime. The combined IM+IN boost was able to enhance plasma anti-HIV-1 IgG, virus neutralization or ADCC titers, milk anti-HIV-1 IgA titers, and antibody epitope diversity (46). Likewise, we found that combined IM+IN boosting increased serum IgG titers, viral neutralization, and ADCC activity in the rabbit model. In fact, combined IM+IN boosting resulted in significantly higher virus neutralizing responses than IM/IM. To prevent functional differences in serum IgG from impacting challenge outcome, we performed NHP studies with the IN/IM+IN regimen as it was the vaccination regimen that induced serum anti-HIV antibody responses most similar to the IM/IM regimen, while also utilizing the highest number of mucosal immunizations to determine if the use of optimized mucosal immunization would influence acquisition of SHIV after repeated low dose rectal challenge.

In NHPs, both the IM/IM and IN/IM+IN regimen induced high serum gp120-specific IgG titers, demonstrating that intranasal vaccines can be included in a vaccination regimen while inducing serum antibody titers similar to those achieved with systemic vaccination regimens. Furthermore, serum gp120-specific IgA was also similar between the vaccine groups after the second and third booster vaccine, with the IN/IM+IN regimen inducing lower IgA responses after administration of the first booster vaccine. Since serum gp120-specific IgA can interfere with protective gp120-specific IgG responses (77), and mucosal vaccination is associated with the production of IgA at mucosal surfaces, the lack of enhancement of serum gp120-specific IgA after the inclusion of intranasal vaccines in the IN/IM+IN regimen demonstrates that mucosal vaccines could potentially be incorporated into HIV vaccination regimens without detrimental IgA-mediated competition with protective IgG for binding to antigens.

While mucosal vaccination is thought to be a better activator of the mucosal immune system than systemic vaccination, the IN/IM+IN vaccine regimen did not induce anti-HIV fecal IgG or IgA greater than those induced by the IM/IM regimen. Additionally, fecal gp120-specific IgG responses were more prevalent and a higher magnitude than fecal gp120-specific IgA responses regardless of the vaccination regimen. There are other examples in the literature where HIV vaccination of NHPs induced antigen-specific mucosal IgG responses but not IgA responses in multiple mucosal samples (60). Our results are consistent with the conclusion that no HIV vaccination regimen developed has induced sustained mucosal IgA responses in human or NHPs(76).

Rabbits were used to optimize intranasal vaccines to induce serum antibody responses similar to those induced by IM vaccination. Rabbits have a nasal cavity anatomy similar to primates(78-80), making them ideal for the development of nasal vaccines. We demonstrated that IM/IM and IN/IM+IN vaccination regimens induced similar serum and mucosal antibody responses in NHPs and rabbits (**Figure 7**). Even though the vaccine regimens evaluated in this study elicited high titer serum antibody responses, and detectable fecal/rectal anti-gp120 IgG titers in both rabbits and NHPs, we did not detect mucosal anti-gp120 IgA in the majority of individuals, although anti-gp120 vaginal IgA responses were observed in 4 of 5 mice nasally immunized with gp120 + M7. Others have also been unable to detect mucosal antigen-specific IgA responses in rabbits after vaccination regimens that resulted in mucosal IgA titers in mice (81). The relatively prolonged intervals between vaccine dosing may also have limited our ability to inducemucosal IgA responses in rabbits and NHPs. For example, we have used nasal immunization schedules of days 0, 7 and 21(16), days 0, 7, 14 and 28(69) or days 0, 21 and 42(21) in mice that induced vaginal antigen-specific IgA while rabbits and NHP were immunized on schedules of 0, 12 and 16 weeks (with a boost at week 24 for the NHP).In addition to its use as an effective animal model for the development of nasally administered vaccines, rabbits are also a useful model for testing Fc-dependent functional antibody responses induced by HIV vaccines (accepted JVI, JVI02119-18R1, Bridging vaccine-induced HIV-1 neutralizing and effector antibody responses in rabbit andrhesus macaque animal models‘). Our work highlights the ability of rabbits to serve as a valuable small animal model system for the development of mucosal HIV vaccination strategies although more studies are needed to better define vaccination regimens that enhance the induction of antigen-specific mucosal IgA responses.

We chose to model our prime/boost regimen off the canarypox prime/ gp120 boost used in the RV144 clinical trial as that prime/boost regimen showed moderate protection from challenge (2). A vaccine regimen that is partially protective should be the most sensitive to any beneficial effects from mucosal vaccines. Unfortunately, no protection from challenge was observed in either vaccine group tested in the present study. Several factors likely played a role in the lack of protection observed. First, others have seen a lack of protection against the same SHIV strain utilized in our study even after several additional boosters of gp120 (51). However, protection was observed when a pentavalent gp120 vaccine was used, and the protective efficacy was correlated to non-neutralizing antibody responses (82). ADCC responses were induced by our prime/boost regimens in rabbits and NHPs, but both the lack of protective efficacy and the ADCC responses observed in our study are similar to what Bradley et al. (51) observed when using a bivalent gp120 vaccine regimen.

The decision to use Addavax, an oil-in-water emulsion, as the adjuvant in our IM administered vaccines instead of alum, which was used in the RV144 trials, may have resulted in lower vaccine efficacy. Vaccari et al., reported that a prime/boost vaccination regimen consisting of priming with the canarypox vector ALVAC and boosting with an adjuvanted gp120 vaccines was effective at preventing SIV acquisition only when the boosting vaccines contained alum (62). The lack of efficacy of booster vaccines adjuvanted with MF-59, another oil-in-water adjuvant, was correlated to mucosal innate lymphoid cells, mucosal V2 antibodies, and RAS activation. Additional studies are needed to determine if mucosal vaccination can overcome any skewing induced by the use of oil-in-water adjuvants for systemic vaccines.

In conclusion, optimized nasal HIV vaccination regimens are capable of inducing high titer, functional serum antibody responses similar to IM vaccination regimens. More work is needed to optimize mucosal immunization for the induction of durable, elevated anti-HIV-1 IgA in mucosal secretions and to determine if mucosal anti-HIV IgA influences protection against mucosal SHIV challenge.

## Acknowledgements

We would like to thank the animal husbandry staff, veterinary technicians, veterinarians, and diagnostic lab personnel at Duke’s Division of Lab Animal Resources for the care they provided our animals and for assistance with animal procedures. We would also like to recognize Moses Wanyonyi and Michael Peace for assistance with rabbit procedures and Carolyn Weinbaum for help obtaining the primates.

For assay and reagent support we would like to thank: Grey Monroe for help performing ELISAs, Raul Louzao and the Immunology & Virology Quality Assessment Center and the Duke Human Vaccine Institute for performing SIV viral loads, Sampa Santra for providing SHIV 1157QNE Y173H, Nicole Rodgers for construction and preparation of MVA viruses, Kevin Saunders and Barton Haynes for providing protein antigens for BAMA, Jamie Peacock and the Duke Human Vaccine Institute Research Protein Production Facility which received funding support from the Collaboration for Aids Vaccine Research Bill and Melinda Gates Foundation(OPP1066832) for C.1086 gp120 protein, and the David Montefiori lab for neutralizing antibody assays which received funding support through the NIH/NIAID Contract # HHSN27201100016C.

